# A mRNA-LNP vaccine against Dengue Virus elicits robust, serotype-specific immunity

**DOI:** 10.1101/2021.01.05.425517

**Authors:** Clayton J. Wollner, Michelle Richner, Mariah A. Hassert, Amelia K. Pinto, James D. Brien, Justin M. Richner

## Abstract

Dengue virus (DENV) is the most common vector-borne viral disease with nearly 400 million worldwide infections each year concentrated in the tropical and subtropical regions of the world. Severe dengue complications are often associated with a secondary heterotypic infection of one of the four circulating serotypes. In this scenario, humoral immune responses targeting cross-reactive, poorly-neutralizing epitopes can lead to increased infectivity of susceptible cells via antibody-dependent enhancement (ADE). In this way, antibodies produced in response to infection or vaccination are capable of contributing to enhanced disease in subsequent infections. Currently, there are no available therapeutics to combat DENV disease, and there is an urgent need for a safe and efficacious vaccine. Here, we developed a nucleotide-modified mRNA vaccine encoding for the membrane and envelope structural proteins from DENV serotype 1 encapsulated into lipid nanoparticles (prM/E mRNA-LNP). Vaccination of mice elicited robust antiviral immune responses comparable to viral infection with high levels of neutralizing antibody titers and antiviral CD4^+^ and CD8^+^ T cells. Immunocompromised AG129 mice vaccinated with the prM/E mRNA-LNP vaccine were protected from a lethal DENV challenge. Vaccination with either a wild-type vaccine, or a vaccine with mutations in the immunodominant fusion-loop epitope, elicited equivalent humoral and cell mediated immune responses. Neutralizing antibodies elicited by the vaccine were sufficient to protect against a lethal challenge. Both vaccine constructs demonstrated serotype specific immunity with minimal serum cross-reactivity and reduced ADE compared to a live DENV1 viral infection.

**IMPORTANCE:** With 400 million worldwide infections each year, dengue is the most common vector-born viral disease. 40% of the world’s population is at risk with dengue experiencing consistent geographic spread over the years. With no therapeutics available and vaccines performing sub optimally, the need for an effective dengue vaccine is urgent. Here we develop and characterize a novel mRNA vaccine encoding for the dengue serotype 1 envelope and premembrane structural proteins that is delivered via a lipid nanoparticle. Our DENV1 prM/E mRNA-LNP vaccine induces neutralizing antibody and cellular immune responses in immunocompetent mice and protects an immunocompromised mouse from a lethal DENV challenge. Existing antibodies against dengue can enhance subsequent infections via antibody-dependent enhancement. Importantly our vaccine only induced serotype specific immune responses and did not induce ADE.

## INTRODUCTION

Dengue virus (DENV) is the most common vector-borne viral disease affecting humans(1–3). Its endemic region now contains 100 countries in Asia, the Pacific, the Americas, and the Middle East(3), with 40% of the world’s population at risk. Disease states during dengue infection manifest as a range of severities; from a self-limiting, febrile illness to more severe cases with life-threatening vascular leakage that can lead to multi-organ failure associated with a viral-driven cytokine storm (4, 5).

DENV is a member of the family *Flaviviridae* of which Zika virus, West Nile virus, yellow fever virus, and Japanese encephalitis virus are also members. It is spread by the arthropod vector *Aedes aegypti* and, to a much lesser extent, *Aedes albopictus* (2, 3). The virus contains a single-stranded, positive-sense RNA genome which codes for a single polypeptide containing three structural proteins; premembrane (prM), envelope (E), and capsid (C), as well as seven nonstructural proteins(6). Dengue virus is categorized into four distinct serotypes, dengue 1-4 (DENV1-4), with amino acid sequence variations of 30-35% across serotypes.

Most countries with endemic dengue are affected by all four serotypes(1). Infection with a single serotype of DENV does not protect against a secondary infection of a heterologous serotype. Instead, primary infection increases an individual’s probability of developing severe clinical symptoms, including shock and death, upon a secondary heterotypic challenge. In this scenario, humoral immune responses after a primary infection produce cross-reactive, non-neutralizing antibodies. These antibodies can bind to infectious virus particles from a secondary, heterotypic challenge and lead to increased infection of cells possessing Fc*γ* receptors via antibody-dependent enhancement (ADE). This poses a challenge for vaccination as a successful vaccine must elicit a neutralizing, long–lasting immune response balanced equally against all four serotypes of DENV.

DENV vaccines that have progressed the furthest in clinical evaluation include CYD-TDV (Dengvaxia, Sanofi-Pasteur), TAK-003 (DENVax, Takeda), and TV003 (NIAID/NIH)(7–11). All three of these vaccines are tetravalent, live-attenuated vaccines which encode for the membrane embedded DENV viral proteins prM and E, in different viral backbones. Other vaccine strategies are in various preclinical stages including recombinant E and subunit vaccines(12–15), purified inactive viruses(16), DNA encoding for prM and E(17, 18), and purified virus-like particles (VLP)(19–21). VLPs, like an infectious viral particle, are comprised of ENV-dimers on the surface resulting in production of particles that share many of the same three-dimensional epitopes as an infectious virus particle(21, 22).

Previously, we developed a mRNA vaccine against the related Zika virus encoding for the viral prM and E proteins(23, 24). This vaccine elicited a robust neutralizing antibody response that protected mice from a lethal Zika viral challenge and prevented vertical transmission of the virus to the fetus. mRNA vaccines have also been shown to provide protective immunity against viral pathogens in non-human primates(25, 26). In this study, we have developed a mRNA vaccine against DENV serotype 1. A construct coding for prM and E proteins was *in vitro* transcribed using the modified nucleotide pseudouridine and resulting mRNA was packaged into lipid nanoparticles (LNP). Following intramuscular injection mRNA-LNPs are taken up into the muscle cells at the site of injection, as well as antigen presenting cells in the draining lymph node(27, 28). Once the cells endocytose the mRNA-LNP, the LNP degrades in the acidified endosome releasing the mRNA into the cytoplasm. The mRNA is then translated into the viral prM-E proteins. The prM-E polyprotein is embedded in the membrane of the ER and cleaved by host protease into the individual viral proteins. The prM and E self-assemble into virus-like particles on the surface of the ER membrane and then the VLP is trafficked through the trans-Golgi network and secreted from the cell. Administration of the DENV1 prM/E mRNA-LNP vaccine elicited neutralizing antibody titers and antiviral specific T cells in wild-type C57BL/6J mice and conferred protection in DENV permissive immunocompromised AG129 mice. Importantly the mRNA-LNP vaccine induced serotype specific immunity with low levels of ADE.

## RESULTS

### Design of DENV1 prM/E Construct and Viral Protein Expression

We designed a construct encoding for the wild-type nucleotide sequence of prM and E proteins from dengue serotype 1 (DENV1) strain 16007 downstream of a Japanese encephalitis virus (JEV) signal peptide. The coding sequence was flanked by a 5’ untranslated region (UTR) previously utilized in other mRNA vaccines(23) and the 3’ UTR from the human hemoglobin subunit alpha 1 mRNA (HBA1) (Figure 1A). The 5’ and 3’ UTRs contribute to translation regulation and mRNA stability essential for optimum protein expression. We *in vitro* transcribed mRNA from a T7 RNA polymerase promoter site upstream of the 5’ UTR. A 5’ cap-1 structure and a 3’ poly-A tail were enzymatically added to produce fully mature messenger RNA that resembles host mRNA. We also generated a separate construct (ΔFL) containing the amino acid substitutions G106R, L107D, and F108A to remove the fusion-loop epitope of the envelope protein. These mutations have been previously characterized and shown to ablate both fusion-loop activity and production of fusion-loop specific antibodies responsible for ADE(29–31).

**Figure 1:**
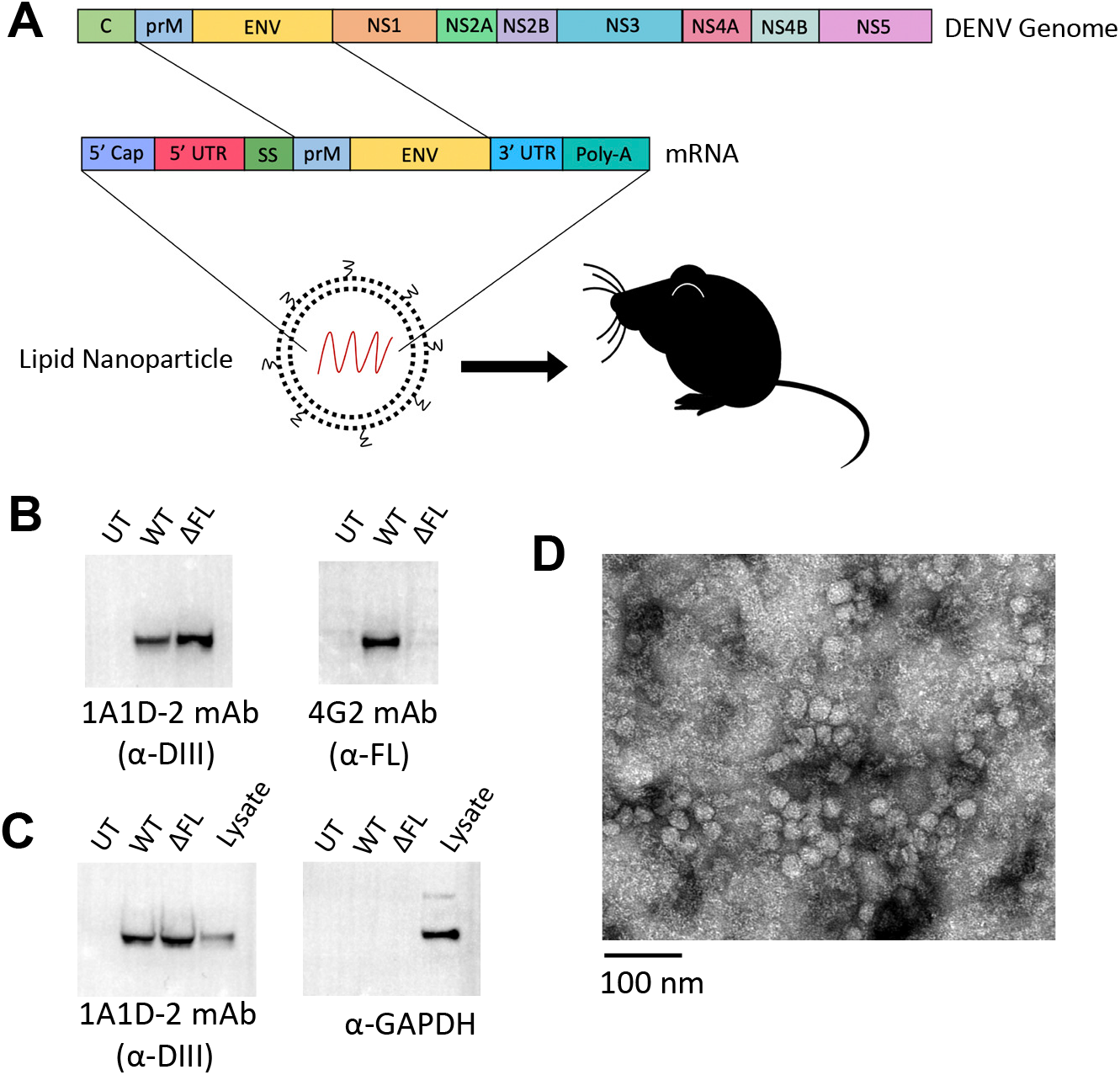
DENV prM/E Vaccine Design and Viral Protein Expression. A) Schematic of the DENV genome and engineered mRNA construct. An mRNA encoding for the prM and ENV viral proteins was engineered with N-terminal signal peptide sequence, 5’ and 3’ untranslated regions (UTR) flanking the coding sequence, a 3’ poly-A tail, and a 5’ cap-1 structure. *In vitro* synthesized mRNA is encapsulated in a lipid nanoparticle for use in *in vitro* and *in vivo* experiments. B) 293T cells were transfected with the *in vitro* transcribed mRNA encoding for the wild-type sequence (WT), or a mutant version with amino acid substitutions in the fusion-loop epitope (ΔFL). Lysate was analyzed by western blot with the domain III specific 1A1D-2 monoclonal antibody and the fusion-loop specific 4G2 monoclonal antibody. C) Supernatant from transfected cells was purified and concentrated through ultracentrifugation and analyzed for VLPs by western blots with the 1A1D-2 monoclonal antibody or anti-GAPDH. Unpurified cell lysate from WT mRNA transfected cells is included as a control. Shown are representative blots. D) Electron microscopy image of VLPs from purified supernatant of transfected 293T cells showing homogenous shape and size of approximately 30nm.

*In vitro* synthesized mRNA was transfected into 293T cells followed by collection of cell lysate and supernatant. We performed immunoblots with the monoclonal antibodies 1A1D-2 and 4G2. 1A1D-2 is specific for domain III of the E protein(32–34) and 4G2 binds to the fusion-loop epitope. Western blots with the monoclonal antibody 1A1D-2 identified a band representing DENV1 E after transfection with both WT and ΔFL constructs (Figure 1B), demonstrating successful viral protein expression. Western blots with 4G2 resulted in a band only in the lysate from wild-type transfected cells, thus revealing successful ablation of the fusion-loop epitope in the ΔFL construct (Figure 1B). Expression of prM and E alone is sufficient to induce the formation and secretion of VLPs(23, 35, 36). To detect secreted VLPs, we purified the supernatant from the transfected cells via ultracentrifugation and analyzed on immunoblots. We detected E protein bands with the 1A1D-2 antibody in the purified supernatant of WT and ΔFL transfected supernatants (Figure 1C), demonstrating that fusion-loop ablation did not affect secretion of VLPs from transfected cells. We did not detect any GAPDH in the purified supernatants verifying that ultracentrifugation removed any cytoplasmic contamination. Particles secreted from transfected cells had similar properties of VLPs with relatively uniform semi-smooth surfaces and diameters of approximately 30nm, as confirmed by electron microscopy (Figure 1D). Together, these results show that *in vitro* synthesized mRNAs can induce viral structural protein expression and secretion of VLPs. Further, mutation of key amino acids within the fusion-loop successfully ablates the antigenic epitope while maintaining protein expression and VLP excretion.

### Optimization of Protein Expression and LNP Delivery

Signal peptides are short N-terminal peptides that traffic proteins through the appropriate processing and secretory pathways within the trans-golgi network. We compared different signal peptides on the DENV1 ΔFL construct to optimize protein expression. We generated five new ΔFL mRNA constructs with the original JEV signal peptide exchanged for signal peptides from either an interleukin-2 (IL2), tissue plasminogen activator (tPA), or gaussia luciferase (GLUC). Additionally, we synthesized two constructs with theoretical signal peptides computationally predicted to elicit robust protein secretion in skeletal muscle cells (SP1 and SP2)(37). Mice are administered the mRNA-LNP vaccine intramuscularly so characterization and optimization of protein expression in muscle cells is key. We transfected differentiated skeletal muscle myoblasts C2C12 cells with the different constructs and blotted for E protein expression with the 1A1D-2 antibody. The TPA signal peptide resulted in the most robust E protein expression (Figure 2A). To ensure that signal peptide modification did not alter VLP secretion and directly compare VLP secretion across the different mRNA constructs, we also analyzed the supernatant of transfected cells via dot blot with the 1A1D-2 mAb. The tPA signal peptide also results in the highest levels of VLP secretion (Figure 2B).

**Figure 2:**
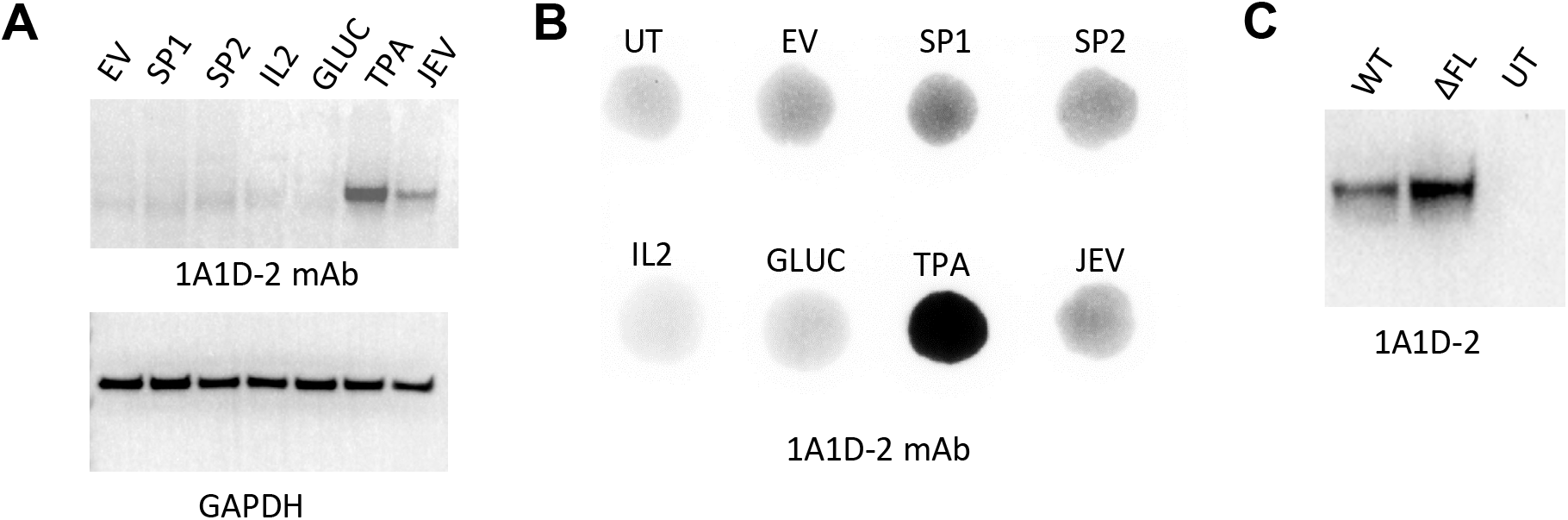
Optimization of Signal Peptide and LNP Delivery. A) Constructs were engineered with alternative signal peptides and *in vitro* transcribed mRNA was transfected into differentiated murine muscle myoblast C2C12 cells. Cell lysate was analyzed by western blot with 1A1D-2 monoclonal antibody or anti-GAPDH antibodies. B) Supernatant of transfected C2C12 cells was analyzed by dot blot with 1A1D-2. C) *In vitro* synthesized WT or ΔFL mRNA was encapsulated into a lipid nanoparticle and administered to C2C12 cells. Lysate was analyzed by western blot with 1A1D-2 antibody. Shown are representative blots.

For *in vivo* administration, mRNA is synthesized with the modified nucleotide, pseudouridine, in place of uridine. This replacement dampens innate immune stimulation and interferon activation which inhibits protein translation(38). *In vitro* synthesized mRNA is further purified and encapsulated in a lipid nanoparticle (LNP). Encapsulation within an LNP shields the mRNA from extracellular RNAses, and ensures efficient delivery into cells(39). LNP are composed of pH sensitive lipids that bind to endogenous apolipoprotein E which facilitates entry. When the mRNA-LNP is endocytosed, the acidic environment of the late endosome initiates degradation of the LNP leading to release of the mRNA to the cytoplasm. We encapsulated mRNA containing the original JEV signal peptide that has been utilized in previous flavivirus mRNA vaccines. We achieved >90% encapsulation efficiency as determined by a ribogreen RNA quantification assay and stored encapsulated mRNA at 4°C for extended periods of time to accommodate a two- or three-shot vaccination schedule. Delivery of nucleotide-modified WT and ΔFL prM/E mRNA-LNPs to C2C12 cells resulted in E protein expression in cell lysate (Figure 2C).

### DENV1 prM/E mRNA Vaccines Elicit Adaptive Immune Responses

Initially, wild-type C57BL/6 mice were vaccinated according to a three-shot vaccination schedule with 10ug of mRNA per dose and serum collections at days 0 (pre-vaccination), day 28 (post-primary), day 42 (post-secondary), and day 56 (post-tertiary) as shown in Figure 3A. We quantified neutralizing antiviral antibody titers in serial dilutions of serum with a focus reduction neutralization test (FRNT) against the homologous DENV1 strain 16007. All mice within each cohort of WT and ΔFL vaccine groups seroconverted with EC50 neutralizing titers (serum concentration at which 50% of the virus is neutralized) maxing out at ~1/200. WT and ΔFL prM/E mRNA-LNP vaccines elicited neutralizing antibody responses after a single dose with secondary and tertiary doses boosting titers (Figure 3B). A third vaccine dose did not significantly enhance the neutralizing antibody titers from that of a second dose (p-value = 0.20, WT). As such, a two-dose, prime-boost vaccination schedule was used in future studies. These data reveal that *in vivo* delivery of a mRNA-LNP vaccine induces a humoral immune response against the exogenous viral protein.

**Figure 3:**
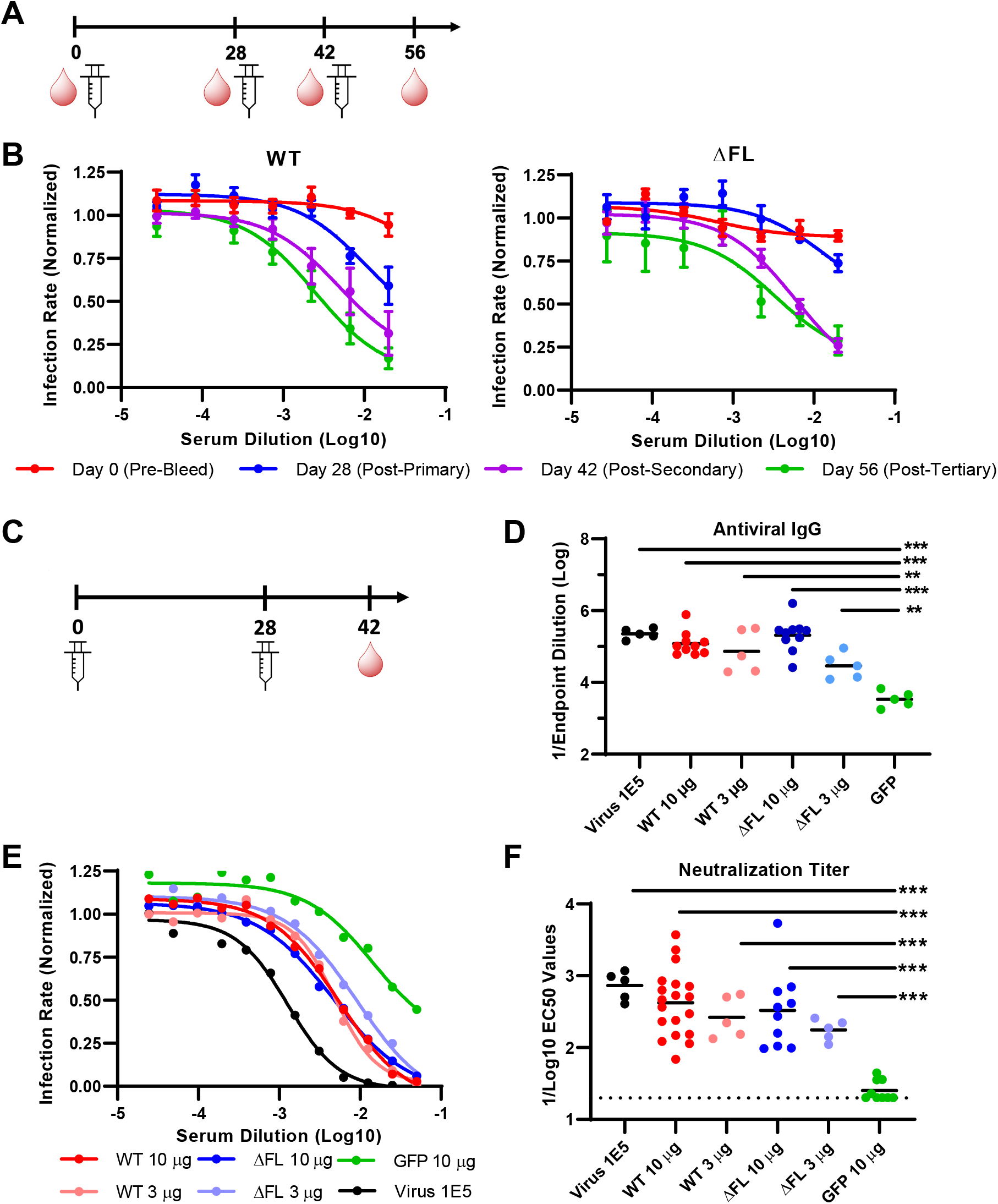
DENV1 prM/E mRNA Vaccines Induce Neutralizing Antibody Response. DENV1 prM/E mRNA-LNP vaccines were administered into 10 week-old C57Bl/6 mice. A.) Mice were administered 10ug of mRNA vaccine in a three-shot schedule and serum collected at the indicated time points. B) Serial dilutions of serum from vaccinated mice were analyzed for neutralization activity with a FRNT assay against DENV1 strain 16007. Neutralization curves at each timepoint are shown for WT vaccine recipients (left) and ΔFL vaccine recipients (right). The average value +/−SEM of five vaccinated mice are shown. C.) Mice were administered a high (10ug) or low (3ug) dose of the mRNA vaccines, or vaccine encoding for GFP. A separate group of mice were infected with wild-type DENV1 following the same schedule. D) Antiviral IgG titers were determined by ELISA assays and the endpoint dilution titer calculated. E) Serum was analyzed by FRNT assays and the normalized percentage infection of each group is plotted as the mean +/−SEM for each serum dilution. N=5 mice per groups of mice infected with virus or receiving 3 ug vaccine doses. N=10 mice per group in mice receiving 10ug doses of the ΔFL, and GFP vaccines. N=15 for mice receiving 10 ug doses of the WT vaccine. F) EC50 values of the neutralization curves from individual mice are shown. Statistical significance of each group compared to the GFP control was determined via unpaired T-test. P-values <0.01 are signified by ** and <0.001 by ***. Statistical comparisons with p-values >0.05 are not shown in this figure.

A separate cohort of mice were administered high or low doses (10μg versus 3μg per injection) of WT or ΔFL prM/E mRNA-LNP vaccine in a prime-boost schedule (Figure 3C). We also included mice infected with live DENV1 (10^5^ FFU DENV1) as a positive control and a mRNA vaccine encoding for GFP (10μg) as a negative control. We quantified the levels of antiviral IgG in the serum isolated from the different vaccine groups via an ELISA assay against purified DENV1 strain 16007. All mice receiving the infectious DENV1, WT mRNA vaccine or ΔFL mRNA vaccine had significantly higher titers than mice receiving the GFP mRNA control vaccine (Fig 3D). No statistical differences were observed between the WT and ΔFL vaccines. Viral-infected mice and mice receiving the high dose of the WT or ΔFL vaccines all had antibody endpoint dilution titers of approximately 1×10^5^. The 3 μg low dose of the vaccine induced antibody titers slightly lower than the higher 10 μg. Serum neutralization titers were determined via FRNT assays (Figure 3E-F) against infectious DENV1 strain 16007. High and low dose of the WT prM/E mRNA vaccine elicited EC50 values of 1/420 and 1/263, respectively, revealing little to no dose dependent response (Figure 3F, p-value = 0.36). Additionally, high and low doses of the ΔFL vaccine resulted in similar EC50 values of 1/329 and 1/175, respectively. These differences were not statistically significant (Figure 3F, p-value = 0.29). The WT and ΔFL vaccinated mice had lower neutralizing titers than the DENV1 virus infected mice (EC50 = 1/729), although these differences were not statistically significant. All vaccines or infections resulted in higher neutralizing values than the GFP vaccinated mice (Figure 3E, p-value <0.001). Neutralizing titers of WT and ΔFL vaccinated mice were very similar, indicating that fusion-loop mutation did not alter humoral immune responses.

To quantify antiviral T cells, splenocytes were harvested from vaccinated mice at day 56 after a tertiary vaccination schedule and stimulated with a pooled 15mer overlapping peptide array for the ENV protein from DENV1 or DENV2 as well as the NS1 protein from DENV1. Stimulated cells were analyzed for intracellular IFNγ by flow cytometry and antiviral IFNγ^+^ T cells were quantified. prM/E mRNA vaccines elicited modest, yet significant antiviral CD4^+^ and CD8^+^ T cell responses specific for the DENV1 ENV protein, with equivalent levels between the WT and ΔFL vaccines (Figure 4 and Supplemental Figure 1). No T cell responses were detected against the homologous DENV2 E protein or the irrelevant DENV1 NS1 protein. Thus, prM/E mRNA-LNP vaccines elicit both humoral and cellular immune responses against DENV1 E protein.

**Figure 4:**
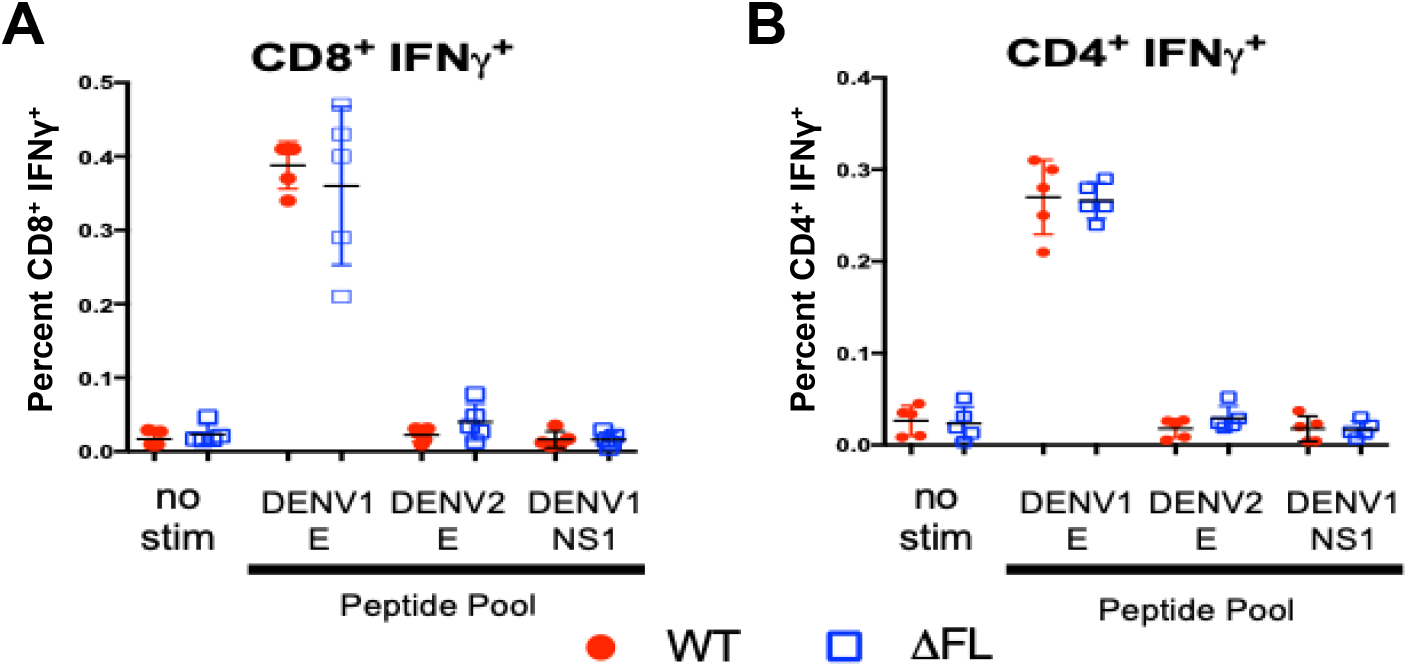
DENV1 prM/E mRNA Vaccines Induces Antiviral CD8^+^ and CD4^+^ T Cells. DENV1 prM/E mRNA-LNP vaccines were administered into 10 week-old C57Bl/6 mice in a three-shot vaccination schedule. Spleens were harvested after the final vaccination dose (day 56) and stimulated with an overlapping peptide array of DENV1 E protein, DENV2 E protein, or DENV1 NS1 protein. Stimulated cells were stained for the intracellular cytokine IFN*γ* and analyzed by flow cytometry. Plotted is the IFNγ^+^ T cells as a percentage of total CD8+ T cells (A) or CD4+ T cells (B). N = 5 mice in the WT and ΔFL vaccinated groups. Representative flow cytometry plots are shown in Supplemental Figure 1.

### DENV1 prM/E mRNA Vaccines Protect against a Lethal Challenge

AG129 mice lack the type I interferon α/β receptor and the type II interferon *γ* receptor, and they are permissible to a lethal DENV challenge(40–42). All serotypes of DENV are capable of replication in AG129 mice with quantifiable viremia, vascular leakage, and increased cytokine levels. Some strains can induce more severe disease states, indicative of severe disease in humans, such as DENV2 D2S20 or DENV1 Western Pacific(40, 42). AG129 mice were vaccinated according to the previously described schedule (Figure 3C) with GFP mRNA-LNP or DENV1 wild-type prM/E mRNA-LNP. Serum was collected from vaccinated mice and analyzed for neutralization titers as previously described. DENV1 prM/E mRNA-LNP vaccination induced EC50 values of greater than 1/3000 (Figure 5A-B). Vaccinated AG129 mice were challenged with 10^6^ FFU DENV1 Western Pacific strain and monitored for 40 days post infection. Mice receiving the GFP mRNA-LNP vaccine lost weight beginning at day 6 and all mice succumbed to viral infection by day 32 post infection. DENV1 prM/E mRNA-LNP vaccinated mice did not show any signs of morbidity or mortality with weight remaining stable post infection and 100% of the mice surviving (Figure 5C-D). These data demonstrate that an DENV1 prM/E mRNA-LNP vaccine protects against a lethal DENV1 challenge in an immunocompromised mouse model.

**Figure 5:**
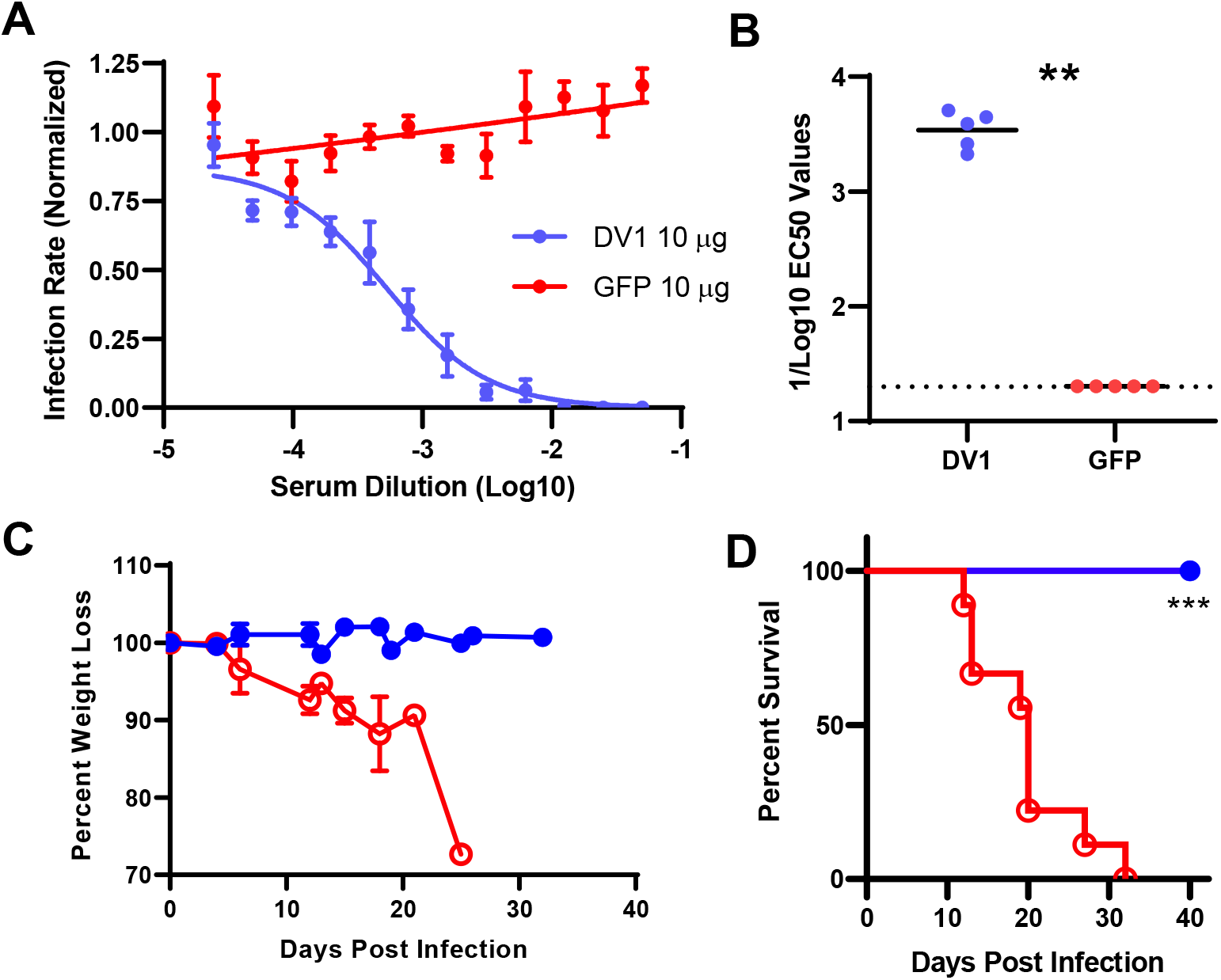
DENV1 prM/E mRNA Vaccines Protect against a Lethal Challenge. 10ug DENV1 prM/E or GFP mRNA-LNP vaccines were administered to AG129 mice in a prime-boost schedule four weeks apart. N = 5 mice per group. A) Serum from vaccinated mice was isolated two-weeks after the boost and analyzed for neutralization by FRNT assay of serially-diluted serum samples. Plotted is the mean +/−SEM from five mice for each dilution. B) EC50 values from each mouse are plotted. The vaccinated mice were then challenged with a lethal dose of DENV1 strain Western Pacific. Mice were monitored for weight (C) and survival (D) post challenge. P-values <0.01 are signified by ** and <0.001 by ***.

The DENV1 prM/E mRNA vaccine elicited both antiviral antibodies and antiviral T cell response. We hypothesized that the antiviral antibodies are sufficient to protect against a lethal challenge. To address this hypothesis, we adoptively transferred pooled serum from WT vaccinated mice into AG129 mice. As controls, a second group of mice received pooled serum from naïve mice and a third group of mice received PBS. One day after adoptive transfer, mice were challenged with 10^6^ FFU DENV1 Western Pacific strain and monitored for 40 days post infection (Figure 6). Seven out of 8 mice that received serum from the vaccinated mice were protected against lethality. Six out of seven mice that received naïve serum lost weight and succumbed to viral lethality post challenge. Thus, antibodies elicited by the DENV1 prM/E mRNA-LNP vaccine are sufficient for protection.

**Figure 6:**
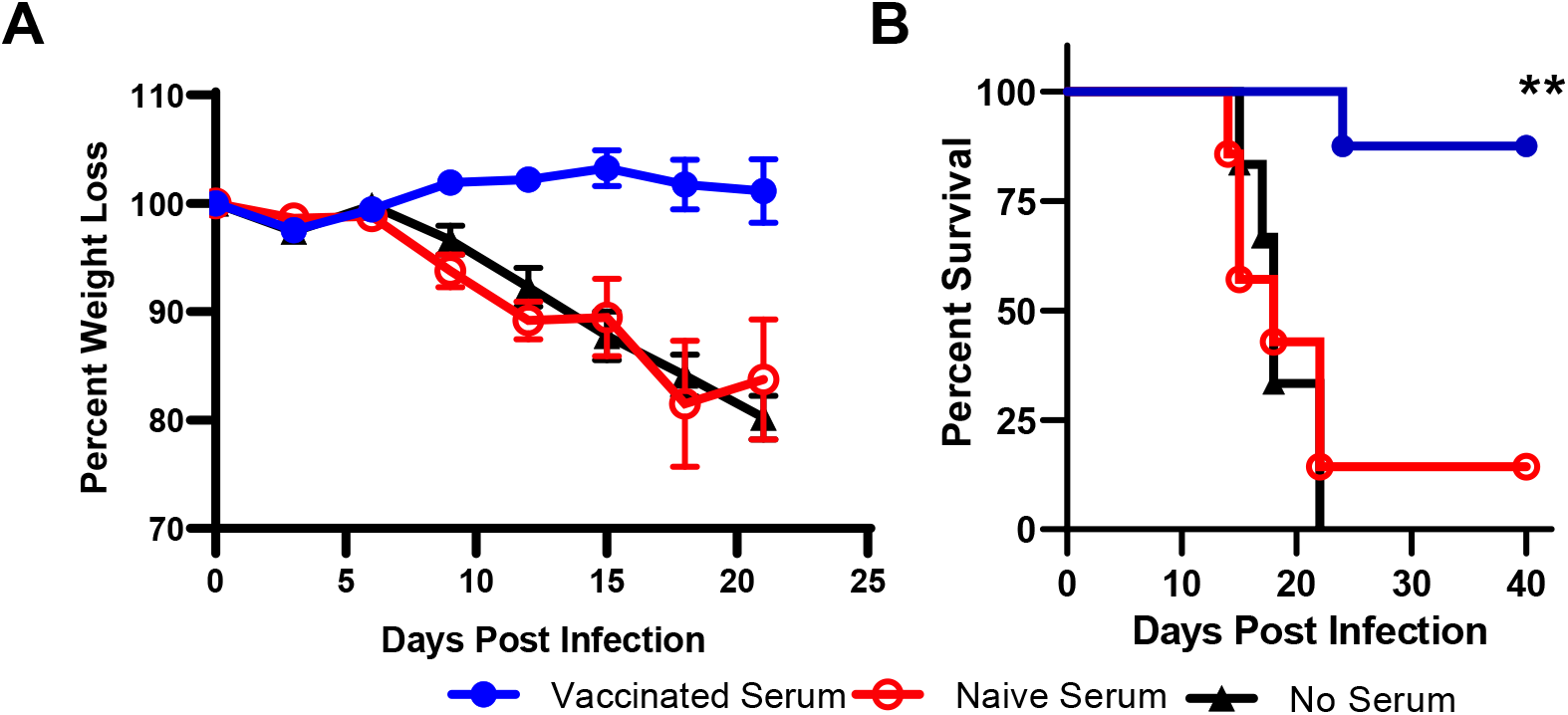
Passive Transfer of Immune Sera Protects Against a Lethal Challenge. Serum from naïve or WT prM/E mRNA vaccinated mice was passively transferred into AG129 mice. One day after transfer, mice were challenged with a lethal dose of DENV1 strain Western Pacific. Mice were monitored for weight (A) and survival (B) post challenge. Survival curves comparing vaccinated and naïve serum recipients were analyzed by log-rank test and p-values <0.01 signified by **.

### DENV1 prM/E mRNA Vaccination Induces Serotype-Specific Humoral Immunity

Infection with DENV will lead to antibodies that cross-react with heterotypic DENV serotypes with the potential to cause ADE. We characterized the cross-reactive immune response in the serum of the prM/E mRNA vaccinated mice. DENV1 vaccines did not elicit neutralizing antibodies against DENV2 (strain New Guinea C) in a FRNT assay (Supplemental Figure 2). We characterized ADE by incubating DENV2 with serial dilutions of serum from vaccinated mice before infecting Fc*γ* receptor-positive K562 cells. Infection was determined via flow cytometry with the monoclonal antibody 1A1D-2 against the viral E protein. The percentage of infected cells was compared to a DENV2 infection in the absence of immune sera (Figure 7A). Serum from DENV1 viral-infected mice significantly enhanced DENV2 infections, even with dilutions as high as 1/6,000. At a serum dilution of 1/100, an 8-fold enhancement was observed. Conversely, serum from mRNA vaccinated mice induced very low levels of DENV2 enhancement (Figure 7A), with only a 1.2-fold enhancement at a 1/100 serum dilution (Figure 7B). The amino acid sequence of the WT prM/E mRNA-LNP vaccine was identical to the sequence of the infecting virus. Surprisingly, WT and ΔFL mRNA vaccines enhanced heterotypic DENV2 to nearly identical values. Similar results were seen with DENV4 (data not shown). As a negative control, serum from naïve mice showed no enhancement at any dilution. To assess the role of ADE on viral replication and egress, we quantified the levels of infectious virus in the supernatant of K562 cells infected with immune-complexed virus. Serum from WT or ΔFL vaccinated mice enhanced viral replication relative to serum from GFP vaccinated mice, however enhancement was significantly lower than serum from viral-infected mice, in agreement with the flow cytometry data (Supplemental Figure 3). These data demonstrate that DENV1 prM/E mRNA vaccines do not induce antibodies which elicit heterotypic enhancement, in contrast to a viral infection.

**Figure 7:**
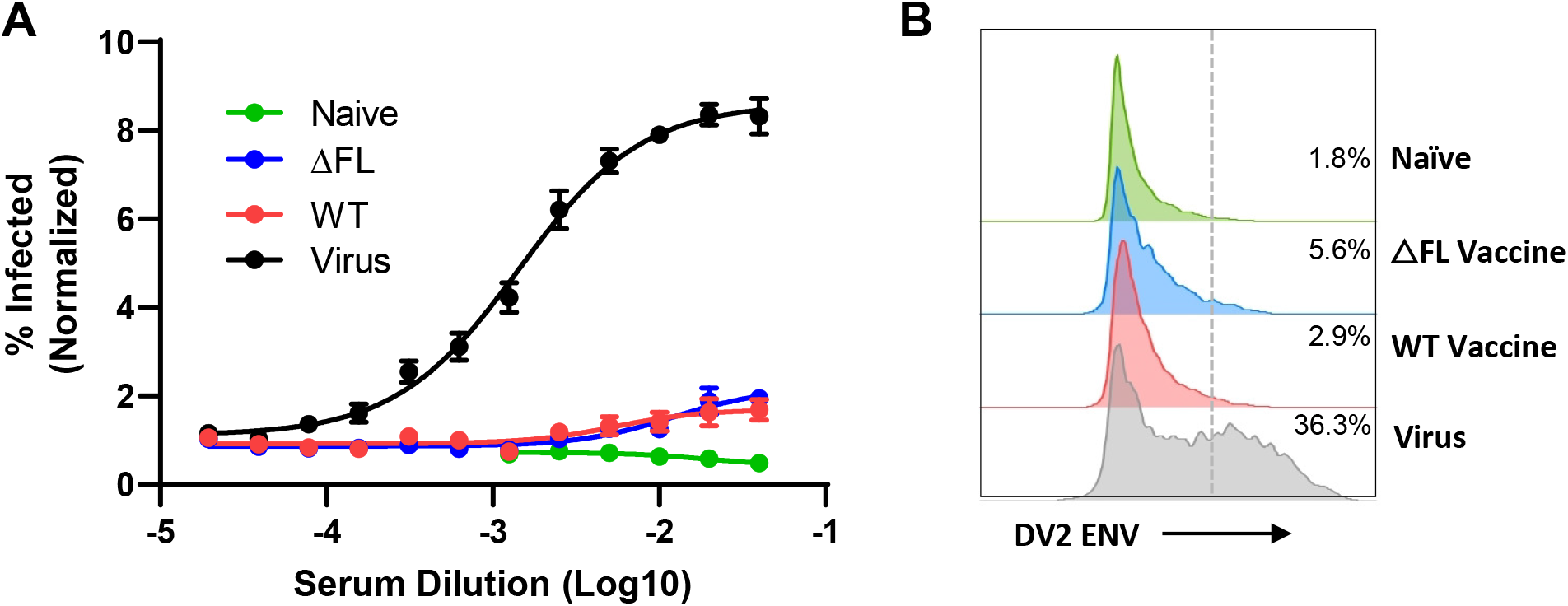
DENV1 prM/E mRNA Vaccination Results in Reduced ADE Levels. Serum from naïve mice, WT prM/E mRNA vaccinated mice, ΔFL prM/E mRNA vaccinated mice, or mice infected with DENV1, two weeks post boost were analyzed for enhancement of DENV2 infection. Serial dilutions of serum were incubated with DENV2 and added to Fc*γ* receptor-positive K562 cells. Fifteen hours later, infected cells were stained for intracellular ENV and quantified by flow cytometry. The percentage of infected cells was normalized to infection in the absence of serum. A.) The fold change in % of cells infected is shown compared to infections in the absence of serum. The average fold enhancement +/−SEM from five mice per group is graphed. B.) A representative flow cytometry histogram of the ENV signal from each different treatment at a 1/100 serum dilution is shown.

## DISCUSSION

Despite a longstanding effort in the field, there still remains an unmet need for a DENV vaccine that elicits robust, balanced immune response against all four serotypes. Here, we developed a vaccine against DENV1 with a modified mRNA encoding for the prM and ENV viral proteins encapsulated in a lipid nanoparticle (LNP). The mRNA-LNP vaccine platform has now been developed for several viruses including rabies virus(26), influenza virus(26), and HIV(43). More recently, mRNA vaccines have been rapidly developed against SARS coronavirus 2 (SARS-CoV-2). Moderna’s mRNA-1273 and Bio-N-Tech mRNA BNT162b1 were the first vaccine candidates to show safety and efficacy in human trials demonstrating the speed of the mRNA platform and its role in emerging infectious diseases(44–46). In the flavivirus field, mRNA vaccines have been developed against Zika virus(23, 24) and Powassan virus(47). These flavivirus vaccines encode for the viral structural proteins which are expressed and lead to the development of neutralizing antibodies against the viral structural proteins. Recently another group published the results of a mRNA vaccine against DENV2(48). Zhang et al developed mRNA vaccines encoding for full length prM-ENV, the soluble portion of ENV, and NS1. Similar to unpublished results from our lab, and contrary to our results with the DENV1 constructs presented herein, Zhang et al observed poor expression of the ENV protein from the prM-ENV DENV2 mRNA construct. Vaccination with the mRNA encoding for the soluble portion of DENV2 ENV (E80) elicited humoral and cell mediated immune responses that protected against a lethal challenge with a homologous serotype of DENV2, similar to our findings with a DENV1 serotype mRNA vaccine. However, the DENV2 E80 mRNA vaccine induced serotype cross-reactive immune responses and high levels of heterologous ADE(48). On the contrary, our mRNA vaccine elicits serotype specific immune responses with low levels of heterotypic ADE.

High level of antigen expression is key for the success of mRNA vaccines. The signal peptide plays a critical role in directing the translated protein into the appropriate locations for processing and secretion. Previous flaviviral mRNA vaccines have included a N-terminal JEV or IgE signal peptide(23, 24, 48). In our study, the tPA signal peptide led to far greater ENV expression and VLP secretion in C2C12 cells compared to other signal peptides, including the JEV signal peptide. All *in vivo* studies here were performed with the original vaccine construct encoding for the JEV signal peptide, but we predict that future vaccine formulations with the tPA signal peptide will lead to greater antigen expression and higher antiviral antibody titers.

The DENV1 mRNA-LNP vaccine elicited humoral and cell mediated immunity following a two-dose vaccination regimen with antibody titers of 1/120,000 and neutralizing titers of 1/420 (WT 10μg). The antiviral antibodies were sufficient for protection. The neutralizing antibody titers reported here are similar to other DENV1 vaccination strategies. Neutralization EC50 titers of approximately 1/100 were achieved by administration of a DNA vaccine encoding for the modified viral structural proteins(29). DENV1 purified-VLP vaccine generated with fusion-loop mutants resulted in neutralization EC50 titers of approximately 1/1,000(31). In phase III human clinical trials, the CYD-TDV (Dengvaxia) vaccine elicited EC50 neutralization titers of approximately 1/60 in seronegative individuals(49), and TAK-003 neutralization titers reached 1/184(50). In phase I human studies TV003 elicited neutralization EC50 titers of 1/63 against DENV1(51).

The neutralizing antibody titers in the AG129 vaccinated mice (EC50 of 1/3,125) were significantly higher than in the C57Bl/6 mice (EC50 of 1/420) following equivalent vaccination schedules (p value <0.001). Likely, the lower neutralization titers in C57Bl/6 mice are due to decreased antigen expression in the presence of an intact type I interferon response. Indeed, previous studies have demonstrated that mRNA vaccines engage RNA-sensing pattern recognition receptors and activate the type I IFN pathway leading to eIF2α phosphorylation and blunted translation of the exogenous transcript(39, 52). Increasing vaccine efficacy could be achieved through lowering the RNA-sensing and IFN response. We have included the pseudouridine modification in our DENV1 prM/E mRNA-LNP vaccine, but further modifications such as 5-methylcytosine could further lower innate immune stimulation and increase antigen expression and associated antibody titers(39).

In humans, CD4^+^ and CD8^+^ T cells predominantly target capsid and NS3, respectively, following a DENV infection(53). Although our vaccine does not encode for these immunodominant T cell epitopes, we detected antiviral CD4^+^ and CD8^+^ T cell responses against the E protein in the vaccinated mice. Intriguingly, the CD4+ and CD8+ T cell responses were not cross-reactive with other DENV serotypes, likely due to the high variability across the different DENV serotypes in the E protein. The overall magnitude of the T cell response from our vaccine was lower than a recently described mRNA vaccine against DENV1 which encoded for the immunodominant HLA epitopes from the nonstructural proteins of DENV(54). Our vaccine was designed to elicit antibodies against the structural proteins to neutralize infectious virus particles as opposed to robust T cell responses. Although we cannot rule out a role for antiviral T cells in a vaccinated host, neutralizing antibodies in serum were sufficient to protect against a lethal homotypic challenge in a passive transfer model. Together these studies demonstrate that mRNA vaccines can be developed to induce both protective T and antibody-dependent immunity against DENV.

Interestingly, the prM/E mRNA vaccines elicited serotype specific antibody responses. Serum from DENV1 virally-infected mice enhanced a DENV2 *in vitro* infection, whereas heterotypic ADE was largely absent with serum from the mRNA vaccinated mice. These observations are surprising given that neutralization titers were similar between the virally-infected and vaccinated mice, and the identical amino acid sequences shared between the WT mRNA vaccine and the infecting DENV1 16007 strain. Further, this suggests that the polyclonal antibody repertoire induced by the mRNA vaccine is inherently different than the polyclonal repertoire induced during a viral infection. In our previous study, mutation of the fusion-loop epitope on the Zika virus mRNA vaccine led to ablation of cross reactive DENV enhancement through ADE(23). Similarly, in previous studies with VLP and DNA based vaccines, mutation of the fusion-loop epitope lowered the prevalence of ADE(29, 55). Unexpectedly, mutation of the fusion-loop epitope in the mRNA vaccine did not alter ADE. These findings suggest that antibodies against the fusion-loop epitope are not dominant in the polyclonal response to our mRNA vaccine. Future efforts will focus on identification of the structural epitopes within the VLP secreted from a viral infection and a mRNA-LNP vaccine.

In this study, we have demonstrated that a mRNA vaccine encoding the prM and E proteins from DENV1 can elicit robust adaptive immune responses and protect against a lethal viral challenge. This study paves the way for future development of mRNA vaccines against the remaining DENV serotypes with the ultimate goal of developing a tetravalent vaccine that will elicit a balanced, protective immune response against all four DENV strains. Current leading vaccination efforts rely on live attenuated virus, yet these vaccines fall short in either their ability to induce a broadly neutralizing antibody response or their ability to avoid ADE. Counter to live attenuated vaccines in which differential replication of the attenuated viruses will dictate antigen dosing *in vivo*, the antigen dose can be carefully modulated with mRNA vaccines to elicit a balanced immune response. Additionally, mRNA vaccines will allow the modification of epitopes which elicit ADE yet are impossible to incorporate into a live attenuated vaccine due to their critical role in viral replication.

## MATERIALS AND METHODS

### Viruses and Cells

DENV serotype 1 strain 16007 and DENV serotype 2 strain New Guinea C was provided by Michael Diamond at Washington University in St. Louis. DENV serotype 4 strain UIS 497 was obtained through BEI Resources (NR-49724), NIAID, NIH as part of the WRCEVA program. Viral stocks were propagated in C6/36 cells and titers determined by a focus-forming assay (FFA). All propagated viral stocks were deep sequenced to confirm viral strain. FFAs were performed to titer viral stocks with monoclonal antibody clone 9.F.10 obtained through Santa Cruz Biotechnology (Cat# SC-70959). Experiments with DENV were conducted under biosafety level 2 (BSL2) containment at the University of Illinois College of Medicine or St. Louis University College of Medicine with institutional Biosafety Committee approval. Vero-E6 cells (Cat# CRL 1586) and K562 cells (Cat# CCL-243) were obtained from American Type Culture Collection (ATCC) and maintained for low passage number following ATCC guidelines. C6/36 cells were provided by the Diamond lab at Washington University in St. Louis, C2C12 cells were obtained from Ahke Heydemann, University of Illinois at Chicago, and 293T cells were obtained from Donna MacDuff at University of Illinois at Chicago.

### Generation of mRNA and mRNA-LNP

Wild-type constructs encoding for dengue serotype 1 strain 16007 prM and Env viral proteins were synthesized by Integrated DNA Technologies (IDT). Constructs contained a T7 promoter site for *in vitro* transcription of mRNA, 5’ UTR and 3’ UTRs, and a Japanese encephalitis virus signal peptide. The sequence of the 5’ and 3’ UTRs were identical to previous publications with a ZIKV mRNA vaccine(23, 24). mRNA was synthesized from linearized DNA with T7 *in vitro* transcription kits from CellScript and following manufacturer’s protocol. Standard mRNA was produced with unmodified nucleotides (Cat# C-MSC11610). RNA to be encapsulated in lipid nanoparticles was generated with pseudouridine in place of uridine with the Incognito mRNA synthesis kit (Cat# C-ICTY110510). 5’ cap-1 structure and 3’ poly-A tail were enzymatically added. mRNA was encapsulated into lipid nanoparticles using the PNI Nanosystems NanoAssemblr Benchtop system. mRNA was dissolved in PNI Formulation Buffer (Cat# NWW0043) and was run through a laminar flow cartridge with GenVoy ILM (Cat# NWW0041) encapsulation lipids at a flow ratio of 3:1 (RNA in PNI Buffer : Genvoy ILM) at total flow rate of 12 mL/min to produce mRNA-LNPs. These mRNA-LNPs were characterized for encapsulation efficiency and mRNA concentration via RiboGreen Assay using Invitrogen’s Quant-iT Ribogreen RNA Assay Kit (Cat# R11490).

### Mouse Experiments

C57BL/6J mice were purchased from Jackson Laboratory and housed in the pathogen free Biomedical Resources Laboratory at University of Illinois College of Medicine. AG129 mice were bred in the animal facilities at Saint Louis University. For vaccinations, mice were injected intramuscularly in the thigh with 50 μl of the indicated amount and type of mRNA-LNP suspended in PBS. Vaccinated C57BL/6J mice were challenged with 1 × 10^5^ FFU of DENV1 strain 16007, retro-orbitally. Vaccinated AG129 mice were challenged with 10^6^ FFU of DENV1 strain West Pac, intravenously. For serum adoptive transfer studies, serum from vaccinated or naïve mice were pooled and then 200μl administered IV into naïve AG129 mice one day prior to challenge with DENV1. The vaccination and viral challenge protocols were approved by the Institutional Animal Care and Use Committee (IACUC) at the University of Illinois College of Medicine (protocol # 18-114) and St. Louis University (assurance # D16-00141).

### *In Vitro* Transfections

293T and C2C12 cells were transfected with mRNA using the Mirus TransIT RNA transfection kit (Cat# MIR 2225) according to manufacturer’s protocol. 293T cells were 60-70% confluent at time of transfection with C2C12 cells being 100% confluent at time of transfection to achieve differentiation into muscle tissue. Supernatant was collected 24 hours post-transfection. To collect lysate, cells were washed with PBS and lysed with RIPA buffer (Millipore-Sigma, Cat# R0278). Lysate and supernatant were centrifuged at 16,000 × g for 10 min at 4°C to remove cell debris. Supernatant from transfected cells was purified using a 20% sucrose cushion and ultracentrifugation at 141,000 × g O/N (16 hours) at 4°C. Purified protein complexes were resuspended in 50μl 1% BSA/PBS for subsequent storage and analysis.

### Viral Protein Detection

For western blot analysis, 10ul of lysate or purified supernatant samples were run on a 4-12% Bis-Tris SDS-PAGE Gel (Invitrogen, Cat #NW04120BOX) with subsequent transfer to 0.45 μm PVDF membrane. Membranes were blocked in TBST (10 nM Tris-HCl, PH 7.5, 150 nM NaCl, and 1% Tween 20) buffer with 5% skim milk. Membranes were blotted with envelope domain III specific 1A1D-2 (1:600) monoclonal antibody (CDC Arbovirus Reference Collection) or envelope fusion-loop specific 4G2 (3.33 mg/ml) (BEI Cat# NR-50327, Novus Biologicals Cat# NBP2-52709FR). Secondary antibody goat anti-Mouse HRP (200 ng/ml) (Invitrogen Cat# A16072) in blocking buffer allowed for detection of dengue viral envelope proteins. Western blots were imaged on ChemiDoc Imagelab system (Bio-Rad).

For dot blot analysis, clarified transfection supernatant was diluted 1/4 in a 20 μl volume of transfer buffer (Life Technologies Cat# NP0006-1) and applied dropwise to a presoaked 0.45 μm PVDF membrane. Sample was allowed to infiltrate membrane through capillary action for no more than one hour (before blot starts to dry). Blots were stained and imaged in the same manner as western blots above.

### Electron Microscopy

One T-75 cell culture flask was seeded with 293T cells at 70-80% confluency the day of transfection. The flask was transfected with 20μg of mRNA encoding WT DENV1 prM and envelope protein using the Mirus TransIT RNA transfection kit (Cat# MIR 2225) according to manufacturer’s protocol. Supernatant was collected 48 hours post-transfection. Supernatant was centrifuged at 16,000 × g for 10 min at 4°C to remove cell debris. 6ml of supernatant was then dialyzed overnight at 4°C in 20,000 MWCO Slide-A-Lyzer Dialysis Cassettes (Cat# 66003) submerged and spinning in PBS. Dialyzed sample was provided to UIC electron microscopy core for imaging using the following parameters. 10-15μl of sample was loaded drop-wise onto a 300-mesh, Formvar/Carbon-coated copper EM grid with excess removed by filter paper via capillary action. One drop 2% Uranyl acetate solution was deposited onto EM grid with excess removed by filter paper via capillary action. Once grid was allowed to dry further, sample was examined via transmission electron microscopy using JEOL JEM-1400F transmission electron microscope, operating at 80 kV. Digital micrographs were acquired using an AMT BioSprint 12M-B CCD Camera and AMT software (Version 701).

### ELISA Assay

Four T-150 cell culture flasks of C6/36 cells were infected with WT DENV1 at an MOI of 0.1. Seven days after infection, 60ml of supernatant was collected and clarified via centrifugation at 3,200 × g for 10 min at 4°C. Supernatant was further purified via 20% sucrose cushion ultracentrifugation at 141,000 × g for 2 hours at 4°C to pellet virus. Virus pellets were resuspended in PBS for a total volume of 5mL. ELISA plates were coated overnight at 4°C with 50μl/well of 1:25 dilution of concentrated viral stock (1E3 FFU/well) in coating buffer (0.1 M sodium carbonate, 0.1 sodium bicarbonate, 0.02% sodium azide at pH 9.6). After coating overnight, plates were incubated with blocking buffer (PBST, 2% BSA, 0.025% Sodium azide) for one hour at 37°C. Plates were then incubated with 50μl of serial dilutions of vaccine and virus enhanced mouse serum at 4°C overnight. Plates were subsequently incubated with goat anti-Mouse HRP secondary antibody (200 ng/ml) (Invitrogen Cat# A16072) in blocking buffer for one hour at room temperature. ELISA plates were developed using 100μl of TMB substrate (Thermo Fisher Cat# 34029). OD 450 reading was measured via BioTek ELISA microplate reader.

### Serum Neutralization Assay

Focus Reduction Neutralization Assays (FRNT) were performed as described previously(23). Briefly, serial dilutions of heat-inactivated serum from vaccinated mice were incubated with 50-70 focus forming units of DENV for one hour at 37°C before infecting a monolayer of Vero cells in a 96 well plate. One hour after infection, cells were overlaid with 1% (w/v) methylcellulose in 2% FBS, 1XMEM. Plates were fixed for 30 minutes with 4% PFA 48 hours after infection. Staining involved 1° antibody 9.F.10 (500 ng/ml) and 2° antibody goat anti-Mouse HRP (200 ng/ml) in PermWash Buffer (0.1% Saponin, 0.1% BSA, in PBS). Treatment with TrueBlue peroxidase substrate (KPL) produced focus forming units that were quantified on the ImmunoSpot® ELISpot plate scanner (Cellular Technology Limited).

### Antiviral T Cell Quantification

Spleens were collected from vaccinated mice and splenocytes collected. An overlapping 15mer peptide library from DENV2 ENV, DENV1 ENV, and DENV NS1 was obtained from BEI Resources, NIAID, NIH, (Catalog #’s NR-507, NR-9241, and NR-2751). Individual peptides were pooled for ex vivo T cell stimulation. Spleens were ground over a 40 μm cell strainer and brought up in Roswell Park Memorial Institute (RPMI) 1640 Medium with 10% FBS, HEPES ((4-(2-hydroxyethyl)-1-piperazineethanesulfonic acid)), and 0.05 mM β-mercaptoethanol. Then, 2×10^6^ cells were plated per well in a round bottom 96-well plate and stimulated for 6 h at 37 °C, 5% CO_2_ in the presence of 10 μg/mL brefeldin A and 10 μg of pooled peptide in 90% DMSO. Following peptide stimulation, cells were washed once with PBS and stained for the following surface markers: α-CD8-PerCP-Cy 5.5 (clone 53–6.7), α-CD3-AF700 (clone 500A2), and α-CD19-BV605 (clone 1D3). Cells were then fixed, permeabilized, and stained for the following intracellular marker: α-IFN-γ-APC (clone B27). The cells were analyzed by flow cytometry using an Attune-NXT.

### ADE Flow Assay

Serial dilutions of heat-inactivated serum from naïve, vaccinated, or viral-infected mice were mixed with DENV2 and incubated for one hour at 37°C. Fc-*γ* receptor (CD32A) positive K562 cells were infected with immuno-complexed virus at an MOI of 1 in a 96 well plate. After a 15-hour incubation, cells were fixed with 4% PFA for 30 minutes and stained for intracellular ENV with 1A1D-2 (1/500) monoclonal antibody and anti-mouse 647-conjugated antibody (2 μg/ml, Invitrogen Cat# A21235).

### ADE Viral Replication Assay

Serum from viral-infected mice and mice receiving the WT, ΔFL, or GFP mRNA vaccine were separately pooled and heat-inactivated. Serum was mixed with DENV2 and incubated for one hour at 37°C at a 1/100 dilution. 10,000 Fc-*γ* receptor (CD32A) positive K562 cells were infected with immuno-complexed virus at an MOI of 1 in a 200μl volume in a 96 well plate. After a 48-hour incubation to allow viral replication and egress, cells were centrifuged to separate cells from supernatant. Viral titers in supernatant were determined via FFA as described above.

## ABBREVIATIONS

DENV: Dengue virus
LNP: Lipid NanoParticle
VLP: Virus Like Particle
prM: premembrane protein
E: envelope protein
FL: fusion-loop
ΔFL: mutant fusion-loop
UT: untransfected
EV: empty vector

## Data Analysis

All data were analyzed with GraphPad Prism software. Statistical significance was determined via unpaired T-tests for comparison of antibody titers, and log-rank tests for comparisons of survival curves. Flow cytometry data was analyzed using FlowJo software (BD Biosciences).

